# Tactile active sensing in insect-plant pollination

**DOI:** 10.1101/2020.06.16.155507

**Authors:** Tanvi Deora, Mahad A. Ahmed, Thomas L. Daniel, Bingni W. Brunton

## Abstract

The interaction between insects and the flowers they pollinate has driven the evolutionary diversity of both insects and flowering plants, two groups with the most numerous species on earth. Insects use vision and olfaction to localize their host plants, but to feed from the flower, they must find the tiny nectary opening, which can be well beyond their visual resolution. When vision is limited, the sense of touch becomes especially crucial, yet very few studies have investigated the role of rapid and precise tactile feedback in successful feeding and pollination interactions. In this paper, we study the remarkable feeding behavior of flying insects that use their proboscis, a flexible mouthpart often longer than their entire body length when unfurled, to expertly explore floral surfaces. Specifically, we observed how the crepuscular hawkmoth *Manduca sexta* interacts with artificial, 3D-printed flowers of varying shapes. We found that moths actively explore the flower for tactile features, systematically sweeping their proboscis from edge to center repeatedly until they locate the nectary. Moreover, naive moths rapidly learn to exploit flowers, and they adopt a tactile search strategy to more directly locate the nectary in as few as three to five consecutive visits. We suggest moths wield their proboscis to extract salient tactile features, such as floral edges and corolla curvature. Our results highlight the proboscis as a unique sensory structure and emphasize the central role of touch in insect-plant pollination interactions.

## Introduction

Insect-plant pollination interactions have shaped the spectacular diversity of both plants and insects. The ability of insects to pollinate flowering plants has been one of the forces that drove the rapid evolutionary diversification of angiosperms (1–3). Further, flowering plants constitute a majority of human agricultural produce, so pollination also serves a core ecological and agricultural service for human populations today (4). Over millions of years, the coevolution of insects and flowering plants has produced a stunning variety of floral specializations. Flowers display species-specific olfactory and visual cues to attract both generalist and specialist insects; insects, in turn, use these cues to find and pollinate their host plants (5–10).

For successful feeding and pollination, insects need not only identify and localize their host flower but also detect the tiny nectary opening on the floral surface. Few studies have focused on the role of rapid and precise mechanosensory feedback in this pollination interaction. For insects like moths and butterflies, this task is particularly difficult, as they hover over the flower to access the nectary with their long, flexible, straw-like mouthpart known as the proboscis. Moreover, crepuscular hawkmoths like *Manduca sexta* are active in dim light conditions at dawn and dusk, when visual feedback is limited by long neural delays (11–14). In addition, the visual resolution of moths at dawn/dusk light levels is on the order of a few centimeters, whereas the nectary opening is no larger than a few millimeters (11, 12). Therefore, insects like moths and butterflies must rely on tactile sensing to successful target the tiny flower nectary opening as they hover over the flowers. The shape and texture of floral surfaces are known to provide both visual and mechanosensory cues in the pollination interaction (15–19), but it remains unknown if insects are actively using and learning tactile cues to find the nectary.

Moths feed with a proboscis, a modified mouthpart that evolved from the two maxillae and is usually held curled up under the head during flight. This mouthpart is heavily muscularized and hydraulically controlled, so that it is both flexible and actively actuated by muscles at its base and along its length (20). The outer tubes are lined with muscles and also carry the trachea and the nerve cord. As a moth approaches a flower, it unfurls the proboscis by pumping body fluid into the two tubes of the proboscis, such that the inner surface of the two tubes zip together to form a central tube called the food canal, which serves as a drinking straw (21). In addition to active control by muscles and hydraulic extension, the proboscis is covered by a vast array of mechanosensory sensillae all along its length and at its base (22, 23). Thus, the proboscis is both a feeding structure and an actively actuated sensory organ whose mechanical properties can be tuned by muscle activation. Perhaps due in part to this complexity, how moths use their proboscis to find the nectary remains poorly understood.

In this paper, we study how moths use their proboscis to actively acquire tactile feedback from the floral surface to feed from the nectary at its center. We also explore how the strategy moths use to extract tactile features changes with learning. We leveraged the natural feeding behavior of hawkmoths to develop a robust, automated behavioral paradigm where tactile cues on flowers are determined by the curvature of their 3D-printed artificial corolla ((18, 19), Figure 1A & B). We tracked the movement of moths and their proboscis tips as they visited and fed repeatedly from these artificial flowers in dim-light conditions, where the tiny opening of the nectary greatly exceeds their visual acuity (Figure 1C). Far from adopting a random search strategy, we found that moths use their proboscis to systematically sweep each flower from edge to center to locate the nectary (Figure 1D). Further, this active exploration improved rapidly, and moths learned a direct strategy to pinpoint the nectary after as few as three to five consecutive visits of the same flower. Touch is a fundamental sensory percept used by all animals to accomplish complex motor behaviors, and it serves a vital role in coordinating the seemingly effortless interactions between one’s body and physical objects in the physical world (24, 25). Understanding how touch shapes these interactions helps us understand a process of great ecological relevance and may also inspire novel haptic technologies.

**Fig. 1.**
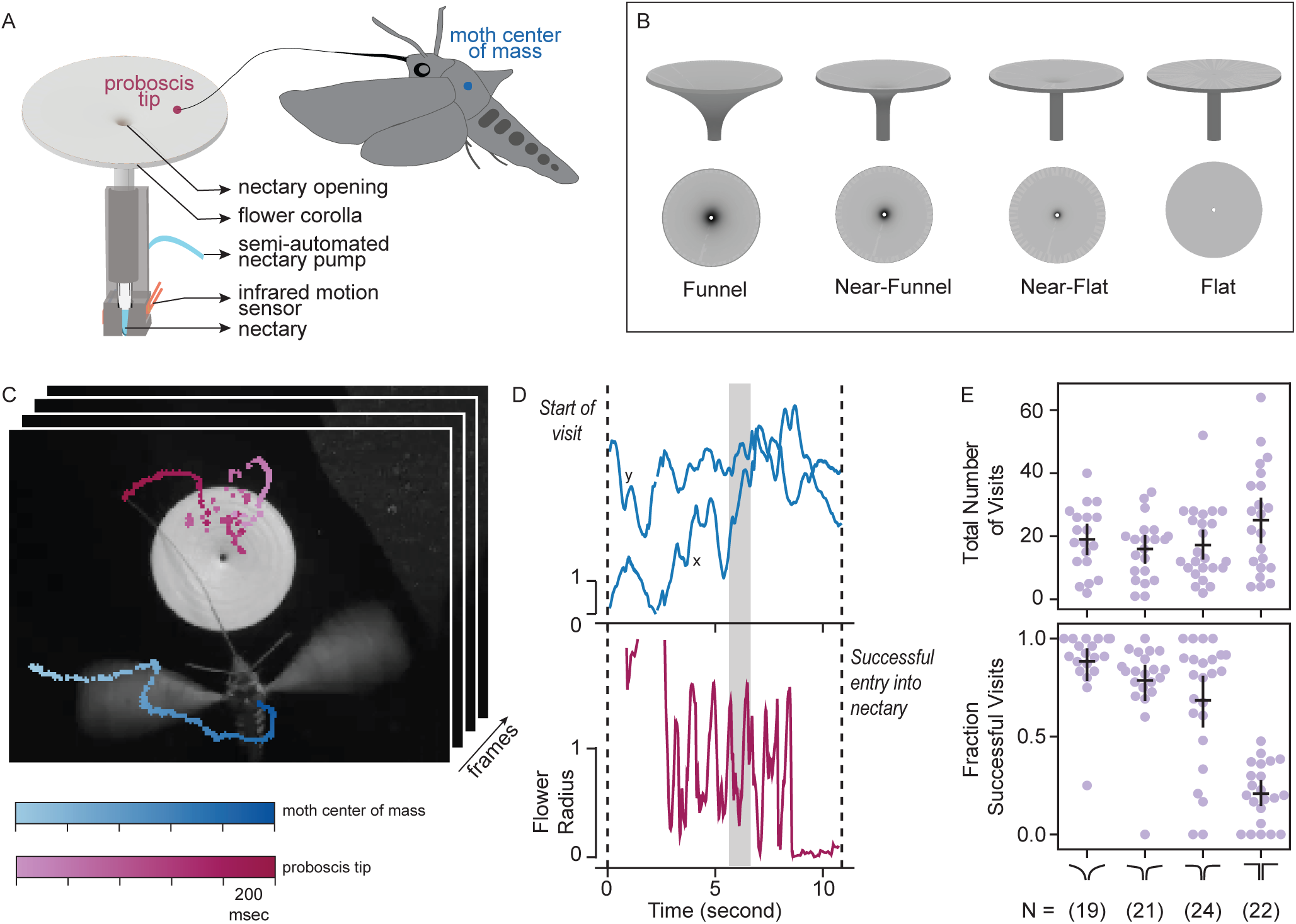
Moths fed from instrumented, 3D-printed artificial flowers as we tracked their body and proboscis tip positions over repeated visits. A) Schematic of a moth visiting an artificial flower. We tracked the proboscis tip and moth’s body using a high speed camera with infrared illumination at 100 frames-per-second. The flower base was instrumented with an infrared motion sensor that triggered when the proboscis reached the nectary. The nectary was also connected to a semi-automated pump that refilled it after each visit (19). B) The four flower shapes in side (top) and overhead (bottom) profiles. All nectaries had the same diameter and height; flowers differed only in the curvature of the their corolla. C) One frame from a video overlaid with moth (blue) and proboscis tip (magenta) tracks. D) Example of tracked trajectories for a single visit, showing the moth position (blue) and proboscis tip (magenta). The gray bar highlights the time window shown in C. The two dashed lines mark exploration time, defined as the start of a visit and the time at which the proboscis enters the nectary. The scale bars represent one flower radius (= 25mm) E) The total number of visits (top) and the fraction of successful visits (bottom) across different floral shapes. The total number of visits are not different across the floral shapes (Kruskal-Wallis H-test *p* = 0.24), however the fraction of successful visits is dependent on floral shape (Kruskal-Wallis H-test *p* = 4.21*e–*09, pairwise Tukey-HSD *p <* 0.05 for funnel/near-flat, funnel/flat, near-funnel/flat and near-flat/flat pairs). Each dot represents an individual moth. The numbers in parenthesis is the total number of moths that interacted at least once with each floral shape.

## Results

We studied the behavior of moths as they explored and learned to feed from flowers of different shapes. In our behavioral paradigm, moths were allowed to feed from 3D-printed flowers in a light controlled chamber while we tracked their centers-of-mass and the tips of their proboscis using a high-speed camera under infrared illumination (Figure 1A & C). All moths were naive to the behavior paradigm and had never fed from any flower (artificial or otherwise) before. Each artificial flower was equipped with a nectary at its base, and this nectary re-filled automatically following successful feeding, so that a single moth could visit the same flower repeatedly. We presented each moth with one of four flower shapes, which differed in the curvature of their corollas (Figure 1B). Each naive moth was tested in a single 30-minute session and with one flower shape; moths visited all flowers with equal likelihood (Figure 1E, Kruskal-Wallis H-test *p* = 0.24, also (18)). Consistent with previous findings, we found that the funnel-shaped flower was the easiest to exploit and the flat flower was the most difficult, as measured by the fraction of visits where the moth accomplished successful feeding over 30 minutes (Figure 1E, Kruskal-Wallis H-test *p* = 4.21*e* –09, pairwise Tukey-HSD *p <* 0.05 for funnel/near-flat, funnel/flat, near-funnel/flat and near-flat/flat pairs).

### Moths learn to feed from different floral shapes over repeated visits

We found that moths quickly learned over repeated visits to the same flower to exploit the curvature of the floral corolla, even when it was very slight, to locate the nectary (Figure 2). We measured how long each moth spent exploring the flower at each visit, defined as the time elapsed between when the moth first comes into the camera view near the flower and when its proboscis reaches the nectary base (Figure 1C, D). In all flower shapes with slight curvature, we found that this exploration time decreased with repeated visits (Figure 2; Kolmogorov–Smirnov (KS) test, *p* = 2.67*e* –11, 1.32*e* –10, 1.45*e* –07 for funnel, near-funnel and near-flat respectively comparing early (highlighted in orange) and late (green) visits). In contrast, for flat flowers, the exploration time did not decrease over repeated visits, suggesting moths did not learn to handle flowers that do not provide surface shape cues to the nectary’s location (*p* = 0.99, KS test).

**Fig. 2.**
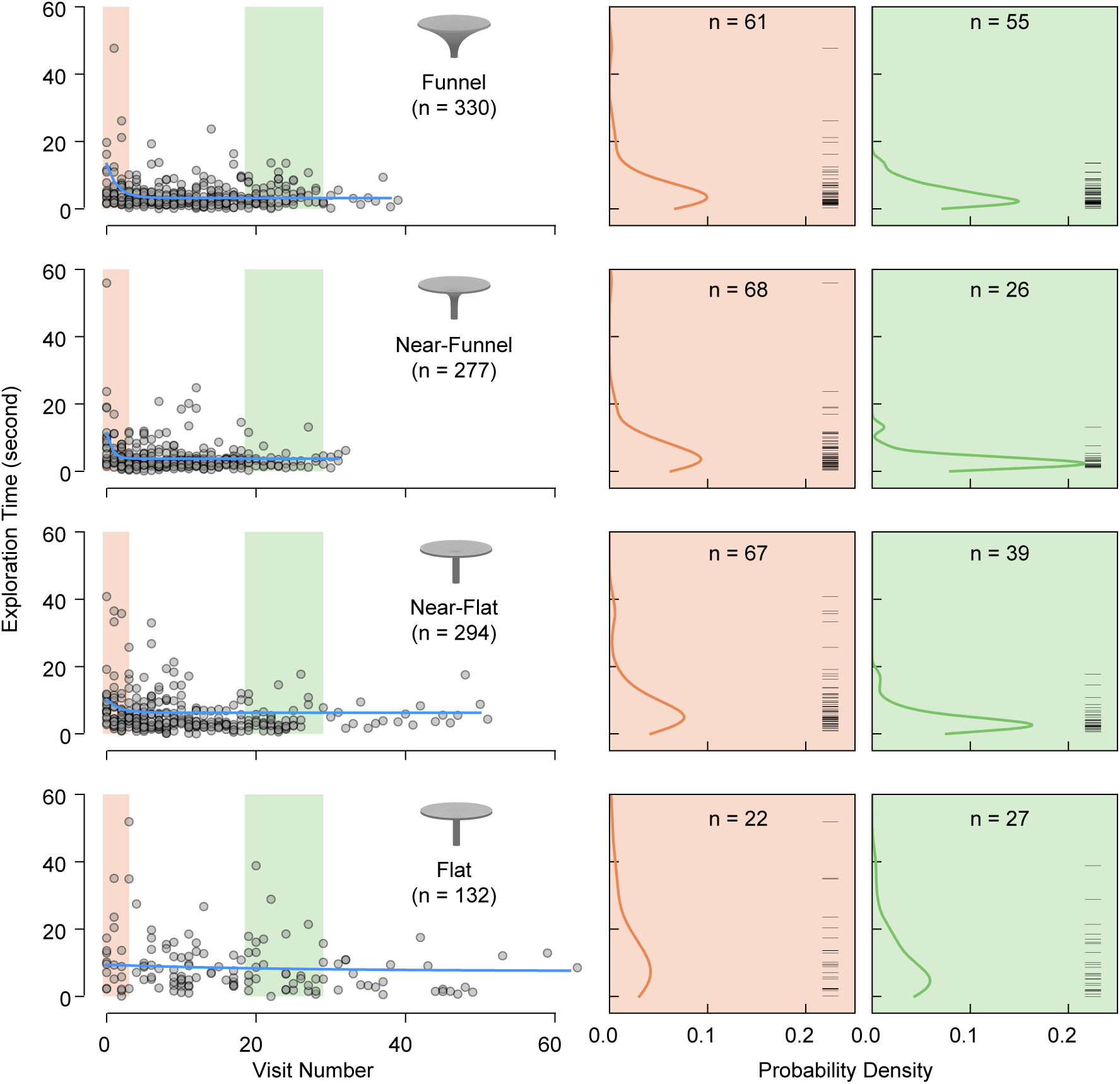
Moths learn to handle flowers in a shape dependent manner. (Left) Learning curves for the four different floral shapes. Each grey dot is the exploration time for a single, successful visit. The blue solid curve shows the exponential fit to this data. In parenthesis, *n* is the number of successful visits pooled across all moths for each flower shape. The exploration times for the early (1st to 3rd visit, in orange) and late (20th to 30th visit, in green) visits pooled and compared on the right. (Right) Probability density estimations of exploration time for early (orange) and late (green) visits. The black ticks on the right in each panel show a raster plot of the raw data used to fit the probability density curves.

Although moths learned after 3–5 visits to handle all three slighted curved flowers, their early exploration times still depended on flower shape. After learning, however, the exploration times did not differ among all shapes except the flat flower (KS test, *p* = 0.97 for funnel/near-funnel, *p* = 0.39 funnel/near-flat, *p* = 0.11 near-funnel/near-flat, and *p* = 1.32*e* –10 for flat/funnel, *p* = 2.67*e* –11 flat/near-funnel, and *p* = 4.05*e* –8 flat/near-flat). Figure 2 shows the aggregated probability densities of all early (first to third, in orange) visits were similar across all flower shapes (KS test *p >* 0.05 for all pairs). However, when we compute how much the distributions diverge from each other using Kullback–Leibler (KL) divergence, we see a flower shape dependent pattern (KL increases as the floral shape diverges: 0.053 funnel/near-funnel < 0.109 funnel/near-flat < 0.228 funnel/flat flowers). Taken together, our data show that moths learn to handle novel flowers that have even slight curvatures within as few as 3–5 visits. Their ability to learn novel floral shapes suggests that moths might be actively extracting cues about floral shape to locate the nectary.

### Moth actively sweep their proboscis to probe flower surfaces

Tracking the tip of the moth proboscis revealed how these mouthparts were used to explore the floral surface, tapping and sweeping, as well as bending the proboscis against the surface (SI Video). To understand the role of the proboscis in flower surface exploration, we trained a neural network to track the trajectory of the proboscis tip in high-speed videos (26). We found that moths explored the floral surface more extensively when attempting to feed from flowers that were more difficult to exploit (Figure 3). These behaviors suggest that moths are extracting mechanical cues during interactions with the flowers.

**Fig. 3.**
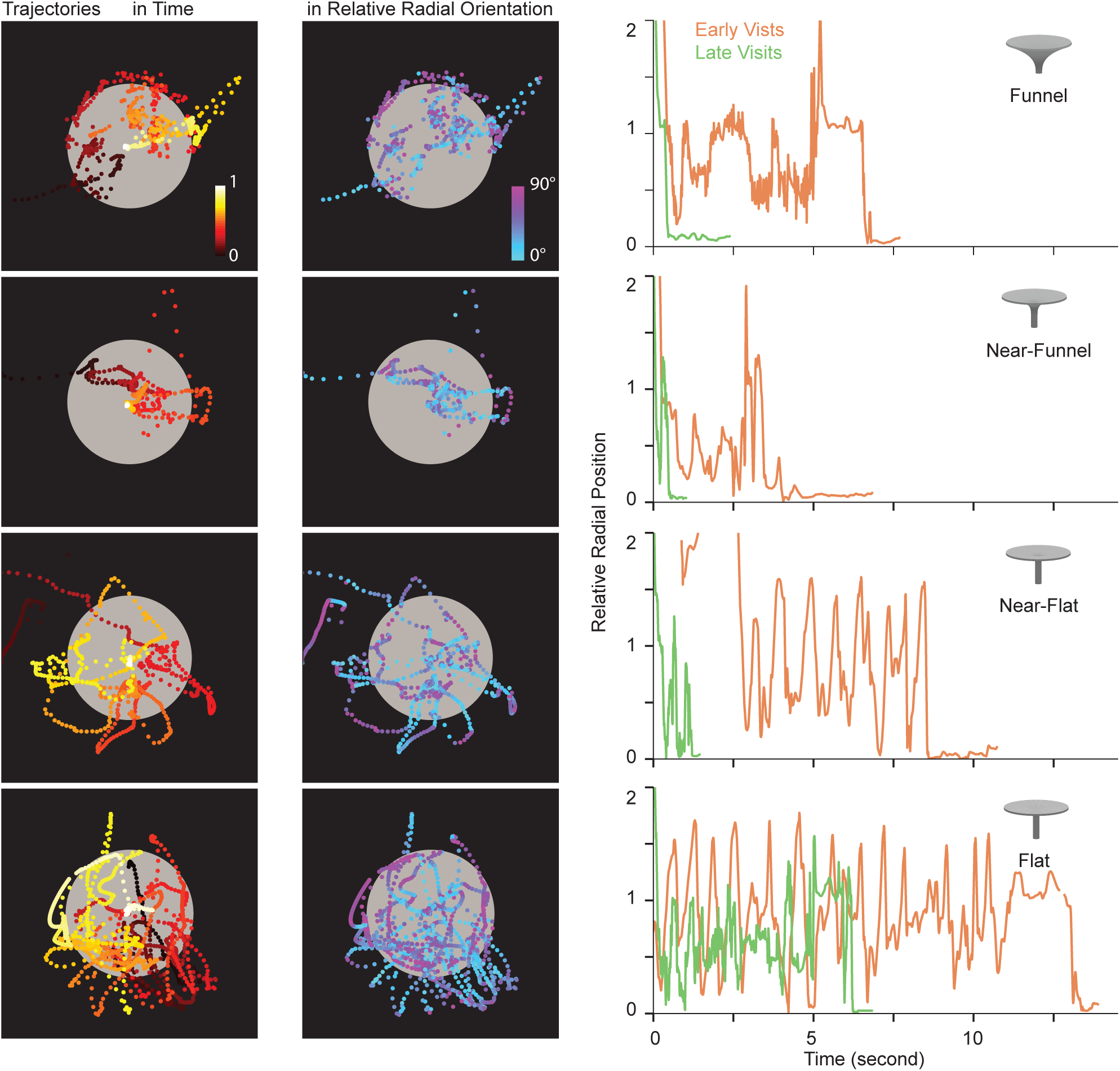
Moths actively explore the floral surface by sweeping their proboscis along the flower. The proboscis tip trajectory for a representative visit for each floral shape, color coded in time (left column), and in relative radial orientation (middle column). (right) The relative radial position of the proboscis tip with time show that for all floral shapes the moths sweep from the edge to the center repeatedly until they find the nectary for early (orange) and later (green) visits.

We next asked if the proboscis tip is moved passively or if the moth manipulated it actively. To disambiguate these possibilities, we examined a few kinematic parameters computed from the proboscis tip tracking. First, we computed the position of the proboscis tip relative to the center of the flower. We found that moths systematically and repeatedly sweep their proboscis between the flower edge and the center as they explore the floral surface (Figure 3). The frequency of sweeps was about 1–2 Hz for all moths and all floral shapes, during earlier and later visits (SF4B, Kruskal-Wallis, *p* = 0.49). Moths found the nectary in fewer sweeps for flowers with even slight curvature as compared to the flat flower (SF4A, Kruskal-Wallis, *p* = 8.65*e* –06). However, the number of peaks did not show systematic, and interpretable changes across visits. For most shapes, as moths learned to handle these flowers, they found the nectary within just a few sweeps. Interestingly, for the most challenging flat flower, moths continued to sweep multiple times for the later visits. These observations are consistent with our previous findings that moths did not learn to feed efficiently from completely flat flowers, despite repeated visits.

Second, we examined the relative radial orientation (RRO) of the proboscis tip with respect to the circular flower corolla. RRO was defined as the angle between the proboscis tip trajectory and the flower’s radial axis. In other words, if the proboscis is sweeping along the radial axis, the RRO would be 0*°*, whereas sweeping perpendicular to the radial axis would have RRO = 90*°* (Figure 3, middle column, colormap blue to purple). Exploring along the radial axis would inform moths about the floral curvature, leading toward the nectary opening at the center. Exploring perpendicular to radial axis would not be informative about the flower surface curvature, except at the edges, where it would inform the moth about the flower’s outer shape. We found that moths explored at RRO of approximately 0*°* along the radial axis on the interior of the flower (Figure 3 and 4). Additionally, for all floral shapes, moths also explored perpendicular to the radial axis along the edge of the flower such that they traced the outer flower profile (Figure 3 and Figure 4).

**Fig. 4.**
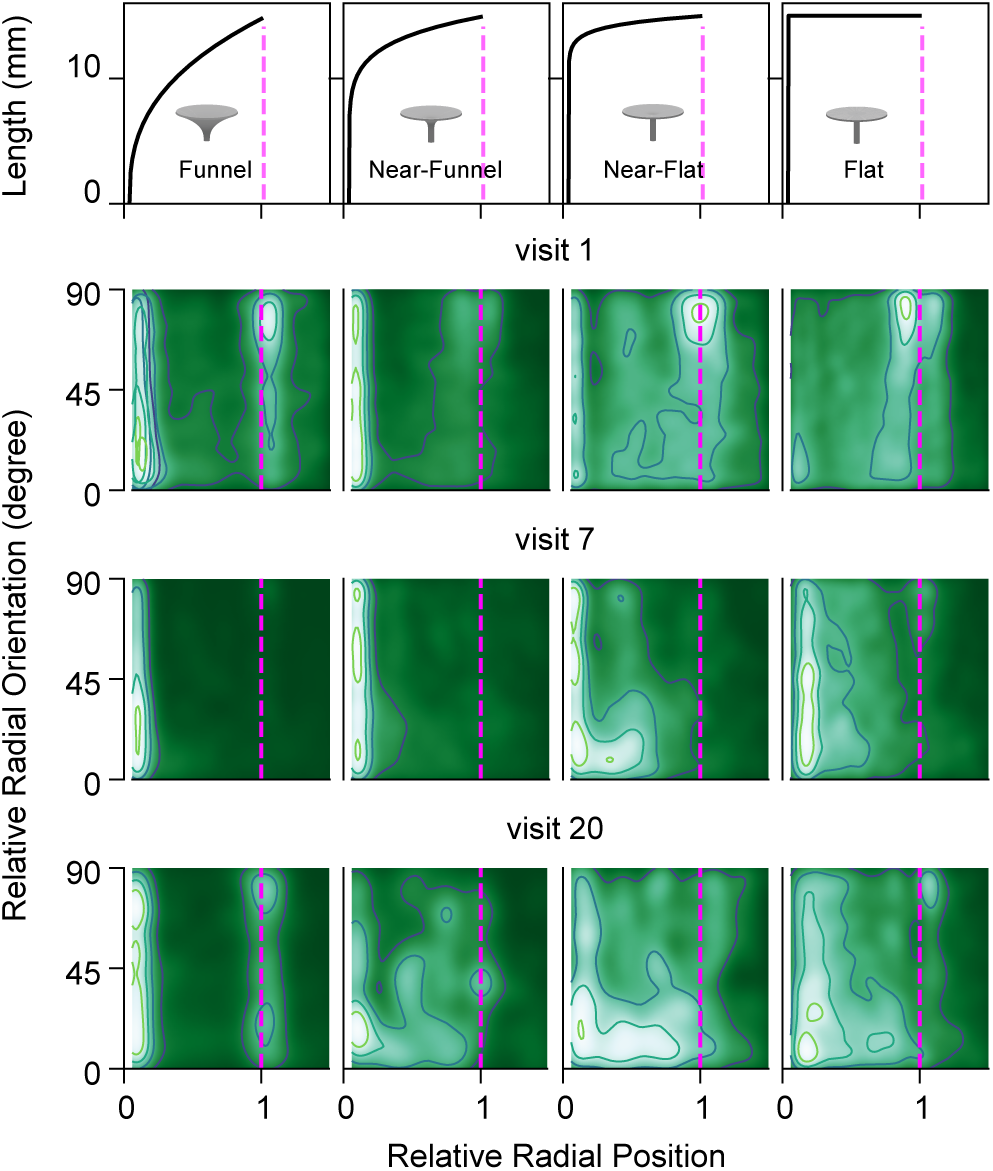
Moths preferentially extract tactile features as they learn to handle novel flower. Heat map of the proboscis tip as a function of relative radial orientation (RRO) and relative radial position for the four floral shapes (along the columns) and over repeated visits (visit 1, 7 and 20; along the rows) pooled across all moths. Peaks in higher probability density (white) shows moths preferentially exploring the edges of the flower (around radial position=1 and RRO=90°). In the interior of the flower, the moths explore at all angles for the first visit. However, over repeated visits, the higher probability density shifts (white) along the radial axis (RRO=0°) for near-funnel and near-flat flowers. (Visit 1 – Funnel: N=6796 frames(16 moths), Near-Funnel:10086 frames(17 moths), Near-Flat:10214 frames(22 moths), Flat:25729 frames(20 moths). Visit 7 – Funnel:2278 frames(13 moths), Near-Funnel:2979 frames(16 moths), Near-Flat:2535 frames(20 moths), Flat:6729 frames(17 moths). Visit 20 – Funnel:1727 frames(8 moths), Near-Funnel:535 frames(5 moths), Near-Flat:1429 frames(9 moths), Flat:5526 frames(13 moths).

### Moths preferentially learn to acquire tactile features

We examined how moths changed their exploration strategies over learning and found that their approaches to gathering tactile cues converged over repeated visits for all floral shapes. Figure 4 visualizes their strategies as probability densities of the proboscis tip trajectories in two dimensions, relative radial orientation (RRO) and relative radial position (for raw data, see SF6). During the first visit, moths spent time exploring along the edge of the flower in all flower shapes, as seen by the high density (in white) around one flower radius with RRO = 90*°*. For the easiest flower shape, across all visits, moths learned to directly find the nectary with only cursory exploration of the floral surface. For more difficult flower shapes, on their first visit, moths explored the interior surface more extensively, at all radial orientations. However, with repeated visits, moths preferentially learned to explore along the radial axis. This radial mode of exploration is evident as the higher density (in white) shifts along RRO = 0*°* for the later visits (7th and 20th visit) for both the near-funnel and near-flat flower. This shift is absent in both the easiest funnel shaped flower and also for the most difficult, flat flower, consistent with the fact that the flat flower has no information about the nectary location along the radial axis.

## Discussion

In summary, our results show that moths use their actively controlled and highly sensed proboscis to explore the three dimensional surface of flowers as they locate the nectary. By high-speed video tracking of proboscis tip trajectories, we characterized how moths systematically sweep their proboscis from edges to centers of flowers as they explored novel floral shapes. Interestingly, as moths learn to exploit the floral corolla curvature over repeated visits, they are able to find the nectary within a few sweeps. Our results suggest that moths use active tactile sensing and learn an efficient strategy over visits to preferentially extract salient mechanical features of the floral edge and curvature.

### Active tactile sensing

A variety of other insects are also known to use tactile feedback to interact with objects in the physical world (27–30). For instance, bees use their antennae and legs to detect the texture of floral surfaces (31, 32). Unlike the smooth lower petal surfaces, the upper (adaxial) petal surfaces are covered in conical shaped epidermal cells (33). These conical cells influence petal color and reflectance, scent release, and petal wettability, in addition to providing a rough, frictional texture surface for use during landing. Moreover, these conical epidermal cells are often arranged in a characteristic spatial pattern that may serve as nectar guides (31). Indeed, bees can be trained to identify specific textures (34). Unlike bees, which land on flowers, hawkmoths like *Maduca sexta* hover as they feed, and thus far floral surface microstructures have not been considered useful for pollination by hovering insects (35). Interestingly, our results suggest that moths do interact with and may leverage textures on the floral surface, and the role of microtextures serving as mechanosensory cues in guiding hovering insect feeding remains to be explored (36)

Touch is a ubiquitous sensory modality across the animal kingdom, and a key feature of tactile sensing is active, and often rhythmic, movement of the sensor to probe and manipulate objects. For example, humans move our fingers to assess the texture of surfaces, and active finger movements lead to improved spatial resolution in touch (37). The use of touch to assess objects and navigate one’s environment has been well studied in diverse organisms, including insects, fish, and rodents (30, 38, 39). In fact, the rat whisker system is among the most well studied examples of active tactile sensing (38, 40). Rats move their whisker bundle rhythmically to feel objects around them, helping them determine the shape and texture of objects, height of obstacles, interact with other con-specifics and navigate through their environment (41). The moth’s proboscis sweeping movements we observed are highly reminiscent of rat whisking.

### Proboscis sensing and mechanics

Although the sweeping motion of hawkmoth proboscis is very similar to the whisking motions of rat whiskers, the sensing and mechanics of the proboscis is entirely different. Rat whiskers are hair shafts of fixed mechanical stiffness, sensed at the base by a single sensory neuron and actuated by muscles at the base (42). Deflections of the whisker shaft can be uniquely mapped to mechanical forces and torques induced at the base of the whisker (43, 44). The sensory neuron at the base can thus faithfully represent contact at the tip by responding to the forces and torques produced at the base. In contrast, the proboscis is hydraulically filled and has muscles not just at its base but along its entire length that actively control its motion, shape, and structural mechanics (23). Further, the proboscis has potential mechanosensors at its base and along its entire length. In its sensing and mechanics, the moth proboscis is closer to a muscular hydrostat like an elephant trunk or an octopus tentacle, except that the proboscis has a stiff cuticular exterior (45). It is interesting to note that although the proboscis is not jointed, it has one relatively fixed point of flexion along its length (SI video). How proboscis mechanics are controlled and how these deformations are sensed are fascinating open questions.

### Multisensory cues in flower exploration

In addition to tactile cues, moths may use feedback from various other sensory modalities to exploit flowers and find the nectary. The visually contrasting grooves on flower surface provide cues that lead to the nectary, and moths have been shown to align their body along the nectar guides (15, 16). Even so, the visual resolution of moths is neither sufficient to resolve the nectary nor provide accurate feedback about proboscis motion. Vision can, however, enable detection of the outer flower contour (46, 47). Thus, in addition to touch, the active movements of proboscis might be also guided by vision, although with limited resolution (12). Indeed, moths handling flat flowers continue to sweep from edge to center despite the lack of curvature cues (Figure 3), even when very few visits were successful (Figure 1).

*Manduca sexta* are crepuscular moths that are active during low light conditions of dusk and dawn, so all of our experiments were conducted at low-light luminance. It is possible that light levels affect the visual control of proboscis motion, and the interaction of vision and touch may be even more crucial for diurnal moths and butterflies in guiding precise proboscis motions. In addition to touch and vision, other sensory cues like humidity gradient over the corolla and gustatory cues on the flower surface might also inform the active movements of proboscis on natural flowers (48, 49).

### Implications of learning and active sensing on pollination and diversity of flowering plants

The mechanistic processes underlying insect-plant pollination are shaped in large part by the insects’ sensory systems and capacity for learning. For instance, bees can identify host plants and learn to associate color or odors with rewards, which influences how they exploit new nectar resources (50). On an evolutionary time scale, these behavioral capabilities may also drive pollination syndromes (51, 52). In other words, insect behavior may drive the evolution of certain floral traits, allowing specific insect species to specialize and exclusively pollinate specific plant species, and hence drive the evolution of new species. Our results show that the hawkmoth, a generalist insect that visits various different kinds of flowering plants, is also an exceptional learner and can learn to handle novel flowers within few visits (Figure 2, also see (17)). However, unlike specialist pollinators such as bees, the success of generalist pollinators like hawkmoths does not necessarily align with the best interest of the plant (19). Plants with flowers that are harder to exploit and more difficult for the insect pollinator to feed from often have greater success in transferring pollen (19). This mismatched interest, coupled with the hawkmoth’s ability to learn and exploit relevant tactile cues in novel flowers, might have profound impact for how they interact with flowers in the wild (53). Thus, understanding the neural basis of sensing and learning in pollinating insects may also shed light on floral diversity and insect forms over evolutionary timescales.

## Supporting information

A moth exploring a slightly curved, near-flat flower with its proboscis during it's first visit.

This file contains additional figures that supplements our results in the main MS.

## Methods

### Moths

We used 2–5 days post-eclosion tobacco hawkmoth, *Manduca sexta* from a colony maintained at the University of Washington. Moths were maintained on a 12:12 hour light-dark cycle. Adults who showed eagerness to feed, assessed in that they flew and hovered in front of a red LED headlamp with their proboscis extended, were selected for experiments. All moths were flower-naive and had never fed prior to experimentation. Moths were dark-adapted for at least 30 minutes before the experiment.

### Behavioral setup

Experiments were conducted in an closed arena (36” × 27” × 36”) with transparent acrylic walls covered by black cardboard. The entire arena was draped with a black cloth to ensure no external light entered the behavioral chamber. All experiments were performed during the active, night period of hawkmoths including dusk and dawn, at about 20–25*°*C. Three viewing windows were cut out of the cardboard to allow video recording and infrared illumination. A high-speed camera (Basler piA640-210gm GigE) was mounted on top of the behavioral chamber and was illuminated using three infrared LED panels. Infrared light is invisible to moths, and hence we used an additional white LED headlamp with a diffuser on one edge of the chamber to simulate dusk/dawn conditions of ∼0.1–1 lux at the flower surface, measured using a light meter (Gossen Mavolux 5032C). We mounted an artificial, 3D-printed flower equipped with micro-sensors (see below) under the camera view. In addition, another funnel shaped distractor flower was placed in the same arena to distract the moths from the rewarded flower between distinct visits. The distractor flower had an empty nectary reserve and moths never received reward for visiting it.

### Artificial flowers

Each moth was presented with one of four flower shapes. The shape of the 3D-printed flower corolla was parameterized by the following equation expressed in cylindrical coordinates (18):

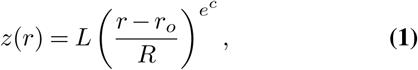

where *z*(*r*) is the longitudinal axis of the flower, and *r* is the radial axis of the corolla from the central z-axis. Each corolla shape is then specified by 4 parameters: *r*_*o*_ = 1mm is the radius of the nectary opening, *R* = 25 mm is radius of the corolla, *L* = 25 mm is the flower’s length, and *c* is a curvature parameter determining the lateral profile of the corolla.

We varied the exponent *c* to generate flowers of different corolla curvature; the funnel, near-funnel, near-flat, and flat flower had *c*= –1, –2, –3 and, ∞ respectively. Flower were 3D printed using white PLA on UPrint printer. Despite high printer resolution, the flower surface had regular concentric grooves from printing layers. We sanded and polished the flower surface using a rotary tool (Dremel 300 series) to provide a smooth surface. The base of 45mm long stalk housed a 200*µ*L PCR tube that served as the nectary reserve (design files can be found at https://github.com/TanviDeora/FlowerDesigns). We filled the nectary with 25*µ*L of 20% sucrose using a semi-automated, custom-built nectary pump at the start of the experiment and also between moth visits ((19), design files can be found at https://github.com/jgsuw/microinjector). Two thin copper wires along the length of the nectary tube detected the presence of nectar. When the nectary was emptied, we prompted the pump to refill the nectary. Additionally, an infra-red transmitter and receiver pair were placed peripheral to the nectary to detect any motion inside the nectary itself. When the moth inserted the proboscis inside the nectary, the light beam became interrupted and the motion was then recorded by a custom written MATLAB script (Arduino and MATLAB codes used can be found at https://github.com/TanviDeora/Arduino-control-codes-for-flight-rig and https://github.com/TanviDeora/MotionVideoCapture respectively). After a successful feeding, if the moth reappeared at the flower in less than 6 seconds, it was not considered as a new visit and moth was not rewarded with nectar.

### Video tracking

To maintain moths in a motivated state, we used a 7-component scent mixture that mimicked the scent of flowers pollinated by hawkmoths ((18); the mixture of volatiles was 0.6% benzaldehyde, 17.6% benzyl alcohol, 1.8% linalool, 24% methyl salicylate, 3% nerol, 9% geraniol, 0.6% methyl benzoate in mineral oil). A few drops of this scent was placed on filter paper and positioned above the rewarded flower on the ceiling of the chamber. We released the moth on one end of the chamber on a raised platform and allowed it to feed repeatedly for a maximum of 30 minutes. The camera captured video at 100 Hz, with 200 *µ*sec exposures, and was time synced with the infrared motion sensor and pump. If a moth failed to interact with the flower within the first 10 minutes, the experiment was concluded. For all flower shapes, we analyzed only those moths that interacted with the flower at least once.

### Analysis of moth Learning

We wrote custom Python scripts (https://github.com/TanviDeora/MothLearning) and used background subtraction to detect the center of mass (moth) and to extract all the instances when the moth appeared in our camera view. Moth and proboscis tip were also tracked by using a trained neural network (DeepLabCut (26)). We classified each instance of moth appearance as a visit if the moth was in view for longer than 1.5 seconds and if the mean likelihood of tracking proboscis tip was greater than 0.4 (based on DeepLabCut tracking). All visits less than 6 seconds apart were merged to be counted as a single visit.

Using the time synced motion sensor data from the nectary, we computed exploration time as the time difference between when the proboscis was detected to have entered the nectary and the start of the visit (Figure 1). All successful visits were used for learning analyses. For each flower type, we fit an exponential decay trend line of the form *y* = *a*_0_ exp(*v/v*_0_)+ *y*_0_ to explain how exploration time changed with visit number *v* for each flower shape. Because the exploration time data is noisy and this exponential decay trend line is sensitive to overfitting to outliers, we estimated *y*_0_ (the asymptotic exploration time after learning) by averaging the last quarter of the data and used the average first-visit exploration time as *a*_0_. The data shown in Figure 2 was then used to fit *v*_0_ by minimizing root mean square error.

We used the exploration times of all moths from their early visits (visit 1–3) and later visits (visit 20–30) to fit estimate probability density functions (PDFs) using a Gaussian kernel density estimator in SciPy. We then used the Kolmogorov-Smirnov (KS) test to evaluate whether these PDFs were different from each other. In addition, we also computed the Kullback-Liebler (KL) divergence to quantify how the PDFs differ from the reference funnel-shaped PDF.

### Proboscis kinematics

We tracked the tip of unfurled proboscis by training a neural network (DeepLabCut (26)). We used 825 manually annotated frames and trained for 1030000 iterations until convergence. To augment the training, a subset of the training data set included frames that were rotated to make the tracking performance rotation invariant.

We wrote custom scripts to smooth the resultant tracking trajectories and to mitigate impact of outliers on our analyses. We computed the distance between the proboscis tip in adjacent frames in manually annotated videos (videos for 6 visits across the 4 floral shapes). Based on this distributions of distances, we estimated the error cutoff to be 24 pixels and used this cutoff to filter DeepLabCut annotation, eliminating all jumps greater than this cutoff. The resulting tracks were smoothed using a median filter (window size 11 time steps) and interpolated with a 3rd order polynomial. Some visits could not be tracked using DeepLabCut and were thus manually tracked.

We computed two kinematic variables from these tracked proboscis trajectories. First, the relative radial distance was computed as the distance of the proboscis tip from the center of the flower normalized to the radius of the flower. Second, the relative radial orientation (RRO) was computed as the angle between the proboscis tip trajectory and radial axis. The angle was wrapped to restrict the range to 0-90*°*. To quantify sweeping behavior, we used points that were less than two radial distance away from center and calculated the number of peaks and frequency of sweeping for all proboscis tip trajectories. We also fit Gaussian kernel estimations to estimate probability density functions (PDFs) and used Kulback-Liebler (KL) divergence to compare the distributions. To analyze relative radial orientation, we ignored parts of trajectory very close to the center (r < 0.06) because the RRO values close to flower center was ambiguous. We plotted tip trajectories distribution in 2D of relative radial distance and relative radial orientation as heat maps and hexbins. We also estimated the PDFs and contours by fitting 2D Gaussian kernels, using a kernel width that was 1.5 times of the bandwidth estimated using Scott’s rule.

## ACKNOWLEDGEMENTS

We would like to thank Joseph Sullivan for help with building the micro-sensed flower and nectary pump, John So for annotating videos and Satpreet Singh and Pierre Karashchuk for help with video tracking. This work was funded by a Human Frontier Science Program Long-Term Fellowship to T.D., Washington Research Foundation Innovation Undergraduate Fellowship in Neuroengineering to M.A.A., grants FA9550-14-1-0398 to T.L.D. and FA9550-19-1-0386 to B.W.B from the Air Force Office of Scientific Research.

## Notes

### Competing Interest Statement

The authors have declared no competing interest.

## Bibliography

1. David Grimaldi. The Co-Radiations of Pollinating Insects and Angiosperms in the Cretaceous. Annals of the Missouri Botanical Garden, 86(2):373–406, 1999.

2. William L. Crepet. Timing in the evolution of derived floral characters: Upper Cretaceous (Turonian) taxa with tricolpate and tricolpate-derived pollen. Review of Palaeobotany and Palynology, 90(3-4):339–359, 1996. ISSN 00346667. doi: 10.1016/0034-6667(95)00091-7.

3. Kathleen M Kay and Risa D Sargent. The Role of Animal Pollination in Plant Speciation : Integrating Ecology, Geography, and Genetics. Annual Review of Ecology, Evolution, and Systematics, 40:637–56, 2009. doi: 10.1146/annurev.ecolsys.110308.120310.

4. Jeff Ollerton, Rachael Winfree, and Sam Tarrant. How many flowering plants are pollinated by animals? Oikos, 120(3):321–326, 2011. ISSN 00301299. doi: 10.1111/j.1600-0706.2010.18644.x.

5. Robert A. Raguso and Mark A. Willis. Synergy between visual and olfactory cues in nectar feeding by wild hawkmoths, Manduca sexta. Animal Behaviour, 69(2):407–418, 2005. ISSN 00033472. doi: 10.1016/j.anbehav.2004.04.015.

6. Joaquín Goyret, Poppy M Markwell, and Robert a Raguso. Context- and scale-dependent effects of floral CO2 on nectar foraging by Manduca sexta. Proceedings of the National Academy of Sciences of the United States of America, 105(12):4565–4570, 2008. ISSN 0027-8424. doi: 10.1073/pnas.0708629105.

7. Jeffrey A. Riffell, Hong Lei, Abrell Leif, and John G. Hildebrand. Neural Basis of a Pollinator’s Buffet: Olfactory Specialization and Learning in Manduca sexta. Science, 339:200–204, 2013.

8. Jeffrey A. Riffell, Hong Lei, Thomas A. Christensen, and John G. Hildebrand. Characterization and Coding of Behaviorally Significant Odor Mixtures. Current Biology, 19(4):335–340, 2009. ISSN 09609822. doi: 10.1016/j.cub.2009.01.041.

9. Almut Kelber. Innate preferences for flower features in the hawkmoth Macroglossum stellatarum. Journal of Experimental Biology, 200(4):827–836, 1997. ISSN 00220949.

10. Almut Kelber. Pattern discrimination in a hawkmoth: Innate preferences, learning performance and ecology. Proceedings of the Royal Society B: Biological Sciences, 269(1509): 2573–2577, 2002. ISSN 14712970. doi: 10.1098/rspb.2002.2201.

11. Jamie C Theobald, Eric J Warrant, and David C O’Carroll. Wide-field motion tuning in nocturnal hawkmoths. Proceedings. Biological sciences / The Royal Society, 277(1683): 853–860, 2010. ISSN 0962-8452. doi: 10.1098/rspb.2009.1677.

12. Anna Lisa Stöckl, David Charles O’Carroll, and Eric James Warrant. Hawkmoth lamina monopolar cells act as dynamic spatial filters to optimize vision at different light levels. Science Advances, 6(16):eaaz8645, 2020. ISSN 23752548. doi: 10.1126/sciadv.aaz8645.

13. Anna Lisa Stöckl, David Charles O’Carroll, and Eric James Warrant. Neural summation in the hawkmoth visual system extends the limits of vision in dim light. Current Biology, 26(6): 821–826, 2016. ISSN 09609822. doi: 10.1016/j.cub.2016.01.030.

14. Simon Sponberg, Jonathan P Dyhr, Robert W Hall, and Thomas L Daniel. Luminance-dependent visual processing enables moth flight in low light. Science, 348(6240):1245–1248, 2015. ISSN 0036-8075. doi: 10.1126/science.aaa3042.

15. Joaquín Goyret and Robert A. Raguso. The role of mechanosensory input in flower handling efficiency and learning by Manduca sexta. The Journal of experimental biology, 209(9): 1585–1593, 2006. ISSN 0022-0949. doi: 10.1242/jeb.02169.

16. Joaquín Goyret. Look and touch: multimodal sensory control of flower inspection movements in the nocturnal hawkmoth Manduca sexta. The Journal of experimental biology, 213 (Pt 21):3676–3682, 2010. ISSN 1477-9145. doi: 10.1242/jeb.045831.

17. Joaquín Goyret and Almut Kelber. Chromatic signals control proboscis movements during hovering flight in the hummingbird hawkmoth Macroglossum stellatarum. PloS one, 7(4): e34629, 2012. ISSN 1932-6203. doi: 10.1371/journal.pone.0034629.

18. Eric O. Campos, Harvey D. Bradshaw, and Thomas L. Daniel. Shape matters: corolla curvature improves nectar discovery in the hawkmoth <i>Manduca sexta</i>. Functional Ecology, 29(4):462–468, 2015. ISSN 02698463. doi: 10.1111/1365-2435.12378.

19. Foen Peng, Eric O. Campos, Joseph Garret Sullivan, Nathan Berry, Bo Bin Song, Thomas L. Daniel, and H. D. Bradshaw. Morphospace exploration reveals divergent fitness optima between plants and pollinators. PLoS ONE, 14(3):1–12, 2019. ISSN 19326203. doi: 10.1371/journal.pone.0213029.

20. G. Wannenmacher and Lutz T. Wasserthal. Contribution of the maxillary muscles to proboscis movement in hawkmoths (Lepidoptera: Sphingidae) – An electrophysiological study. Journal of Insect Physiology, 49(8):765–776, 2003. ISSN 00221910. doi: 10.1016/S0022-1910(03)00113-6.

21. Chengqi Zhang, Peter H. Adler, Daria Monaenkova, Taras Andrukh, Suellen Pometto, Charles E. Beard, and Konstantin G. Kornev. Self-assembly of the butterfly proboscis: The role of capillary forces. Journal of the Royal Society Interface, 15(144), 2018. ISSN 17425662. doi: 10.1098/rsif.2018.0229.

22. Harald W Krenn. Functional morphology and movements of the proboscis of Lepidoptera (Insecta). Zoomorphology, 110(2):105–114, 1990. ISSN 0720-213X. doi: 10.1007/BF01632816.

23. Harald W Krenn. Feeding mechanisms of adult Lepidoptera: structure, function, and evolution of the mouthparts. Annual review of entomology, 55(36):307–327, 2010. ISSN 0066-4170. doi: 10.1146/annurev-ento-112408-085338.

24. Hannes P. Saal and Sliman J. Bensmaia. Touch is a team effort: Interplay of submodalities in cutaneous sensibility. Trends in Neurosciences, 37(12):689–697, 2014. ISSN 1878108X. doi: 10.1016/j.tins.2014.08.012.

25. Mathew E Diamond and Ehsan Arabzadeh. Progress in Neurobiology Whisker sensory system – From receptor to decision. Progress in Neurobiology, 2012. ISSN 0301-0082. doi: 10.1016/j.pneurobio.2012.05.013.

26. Alexander Mathis, Pranav Mamidanna, Kevin M. Cury, Taiga Abe, Venkatesh N. Murthy, Mackenzie Weygandt Mathis, and Matthias Bethge. DeepLabCut: markerless pose estimation of user-defined body parts with deep learning. Nature Neuroscience, 21(9):1281–1289, 2018. ISSN 15461726. doi: 10.1038/s41593-018-0209-y.

27. J. Erber, S. Kierzek, E. Sander, and K. Grandy. Tactile learning in the honeybee. Journal of Comparative Physiology – A Sensory, Neural, and Behavioral Physiology, 183(6):737–744, 1998. ISSN 03407594. doi: 10.1007/s003590050296.

28. Christoph Schütz and Volker Dürr. Active tactile exploration for adaptive locomotion in the stick insect. Philosophical Transactions of the Royal Society B: Biological Sciences, 366 (1581):2996–3005, 2011. ISSN 14712970. doi: 10.1098/rstb.2011.0126.

29. C. M. Comer, L. Parks, M. B. Halvorsen, and A. Breese-Terteling. The antennal system and cockroach evasive behavior. II. Stimulus identification and localization are separable antennal functions. Journal of Comparative Physiology A: Neuroethology, Sensory, Neural, and Behavioral Physiology, 189(2):97–103, 2003. ISSN 03407594. doi: 10.1007/s00359-002-0384-9.

30. J. M. Camhi and E. N. Johnson. High-frequency steering maneuvers mediated by tactile cues: Antennal wall-following in the cockroach. Journal of Experimental Biology, 202(5): 631–643, 1999. ISSN 00220949.

31. P. G. Kevan and M. A. Lane. Flower petal microtexture is a tactile cue for bees. Proceedings of the National Academy of Sciences, 82(14):4750–4752, 1985. ISSN 0027-8424. doi: 10.1073/pnas.82.14.4750.

32. Cwyn Solvi, Selene Gutierrez Al-Khudhairy, and Lars Chittka. Bumble bees display cross-modal object recognition between visual and tactile senses. Science, 367(6480):910–912, 2020. ISSN 10959203. doi: 10.1126/science.aay8064.

33. Heather M. Whitney, K. M.Veronica Bennett, Matthew Dorling, Lucy Sandbach, David Prince, Lars Chittka, and Beverley J. Glover. Why do so many petals have conical epidermal cells? Annals of Botany, 108(4):609–616, 2011. ISSN 03057364. doi: 10.1093/aob/mcr065.

34. Heather M. Whitney, Lars Chittka, Toby J.A. Bruce, and Beverley J. Glover. Conical Epidermal Cells Allow Bees to Grip Flowers and Increase Foraging Efficiency. Current Biology, 19 (11):948–953, 2009. ISSN 09609822. doi: 10.1016/j.cub.2009.04.051.

35. M. Kraaij and C. J. Kooi. Surprising absence of association between flower surface microstructure and pollination system. Plant Biology, pages 1–7, 2019. ISSN 1435-8603. doi: 10.1111/plb.13071.

36. Tobias Policha, Aleah Davis, Melinda Barnadas, Bryn T.M. Dentinger, Robert A. Raguso, and Bitty A. Roy. Disentangling visual and olfactory signals in mushroom-mimicking Dracula orchids using realistic three-dimensional printed flowers. New Phytologist, 210(3):1058–1071, 2016. ISSN 14698137. doi: 10.1111/nph.13855.

37. Lisa Skedung, Martin Arvidsson, Jun Young Chung, Christopher M. Stafford, Birgitta Berglund, and Mark W. Rutland. Feeling small: Exploring the tactile perception limits. Scientific Reports, 3, 2013. ISSN 20452322. doi: 10.1038/srep02617.

38. Mathew E Diamond, Moritz Von Heimendahl, and Per Magne Knutsen. ‘Where’ and ‘what’ in the whisker sensorimotor system. Nature Reviews Neuroscience, 9:601–613, 2008. doi: 10.1038/nrn2411.

39. J. Erber, B. Pribbenow, A. Bauer, and P. Kloppenburg. Antennal reflexes in the honeybee: tools for studying the nervous system. Apidologie, 24(3):283–296, 1993. ISSN 0044-8435. doi: 10.1051/apido:19930308.

40. Nicholas E Bush, Sara A Solla, and Mitra J Z Hartmann. Whisking mechanics and active sensing. Current Opinion in Neurobiology, 40:178–188, 2016. ISSN 0959-4388. doi: 10.1016/j.conb.2016.08.001.

41. David Kleinfeld, Ehud Ahissar, and Mathew E Diamond. Active sensation : insights from the rodent vibrissa sensorimotor system. Current Opinion in Neurobiology, 16:435–444, 2006. doi: 10.1016/j.conb.2006.06.009.

42. Rune W Berg and David Kleinfeld. Rhythmic Whisking by Rat : Retraction as Well as Protraction of the Vibrissae Is Under Active Muscular Control. Journal of Neurophysiology, 89:104–117, 2003.

43. Nicholas E Bush, Christopher L Schroeder, Jennifer A Hobbs, Anne E T Yang, Lucie A Huet, Sara A Solla, and Mitra J Z Hartmann. Decoupling kinematics and mechanics reveals coding properties of trigeminal ganglion neurons in the rat vibrissal system. eLIFE, pages 1–23, 2016. doi: 10.7554/eLife.13969.

44. Lucie A Huet, Christopher L Schroeder, and Mitra J Z Hartmann. Tactile signals transmitted by the vibrissa during active whisking behavior. Journal of neurophysiology, 113:3511–3518, 2015. doi: 10.1152/jn.00011.2015.

45. William M. Kier and Kathleen K. Smith. Tongues, tentacles and trunks: The biomechanics in muscular hydrostats, 1985.

46. Eatai Roth, Robert W Hall, Thomas L Daniel, and Simon Sponberg. Integration of parallel mechanosensory and visual pathways resolved through sensory conflict. Proceedings of the National Academy of Sciences of the United States of America, 113(45):12832–12837, 2016. ISSN 1091-6490. doi: 10.1073/pnas.1522419113.

47. Anna L. Stöckl, Klara Kihlström, Steven Chandler, and Simon Sponberg. Comparative system identification of flower tracking performance in three hawkmoth species reveals adaptations for dim light vision. Philosophical Transactions of the Royal Society B: Biological Sciences, 372(1717), 2017. ISSN 14712970. doi: 10.1098/rstb.2016.0078.

48. Martin Von Arx, Joaquín Goyret, Goggy Davidowitz, and Robert A. Raguso. Floral humidity as a reliable sensory cue for profitability assessment by nectar-foraging hawkmoths. Proceedings of the National Academy of Sciences of the United States of America, 109(24): 9471–9476, 2012. ISSN 00278424. doi: 10.1073/pnas.1121624109.

49. Alexander Haverkamp, Felipe Yon, Ian W. Keesey, Christine Mißbach, Christopher Koenig, Bill S. Hansson, Ian T. Baldwin, Markus Knaden, and Danny Kessler. Hawkmoths evaluate scenting flowers with the tip of their proboscis. eLife, 5(MAY2016):1–12, 2016. ISSN 2050084X. doi: 10.7554/eLife.15039.

50. Lars Chittka and Nigel E. Raine. Recognition of flowers by pollinators. Current Opinion in Plant Biology, 9(4):428–435, 2006. ISSN 13695266. doi: 10.1016/j.pbi.2006.05.002.

51. Steven D Johnson. Phylogenetic evidence for pollinator- driven diversification of angiosperms. Trends in Ecology & Evolution, 27(6):353–361, 2012. doi: 10.1016/j.tree.2012.02.002.

52. Charles B. Fenster, W. Scott Armbruster, Paul Wilson, Michele R. Dudash, and James D. Thomson. Pollination syndromes and floral specialization. Annual Review of Ecology, Evolution, and Systematics, 35:375–403, 2004. ISSN 00664162. doi: 10.1146/annurev.ecolsys.34.011802.132347.

53. Richard E Walton, Carl D Sayer, Helen Bennion, and Jan C Axmacher. Nocturnal pollinators strongly contribute to pollen transport of wild flowers in an agricultural landscape. Biology letters, 16(20190877), 2020.

