## Supplementary material for "Tactile active sensing in insect-plant pollination": This file contains additional figures that supplements our results in the main MS.

### The role of tactile active sensing in insect-plant pollination - Supplementary information

This section supplements the main text, providing additional figures to support our results.

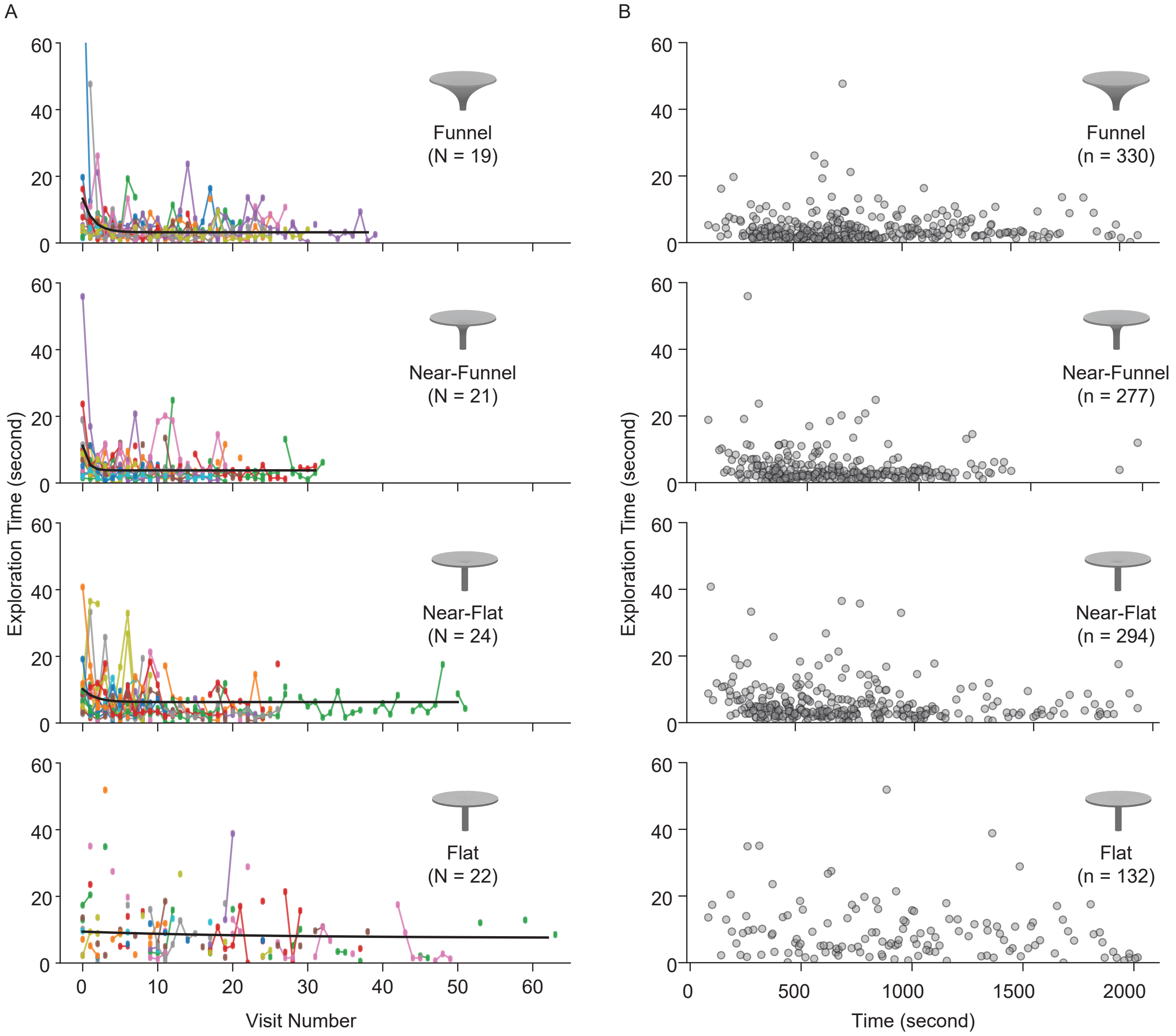


**Fig. 1.** (A) Learning curves for individual moths across all floral shapes are plotted as versus visit number (panel A) and time (panel B). Each color in panel A is for a separate individual. The black solid line is the exponential fit to the data pooled across all moths as shown in Fig 2 in main MS. Each dot in panel B represents a single successful visit.

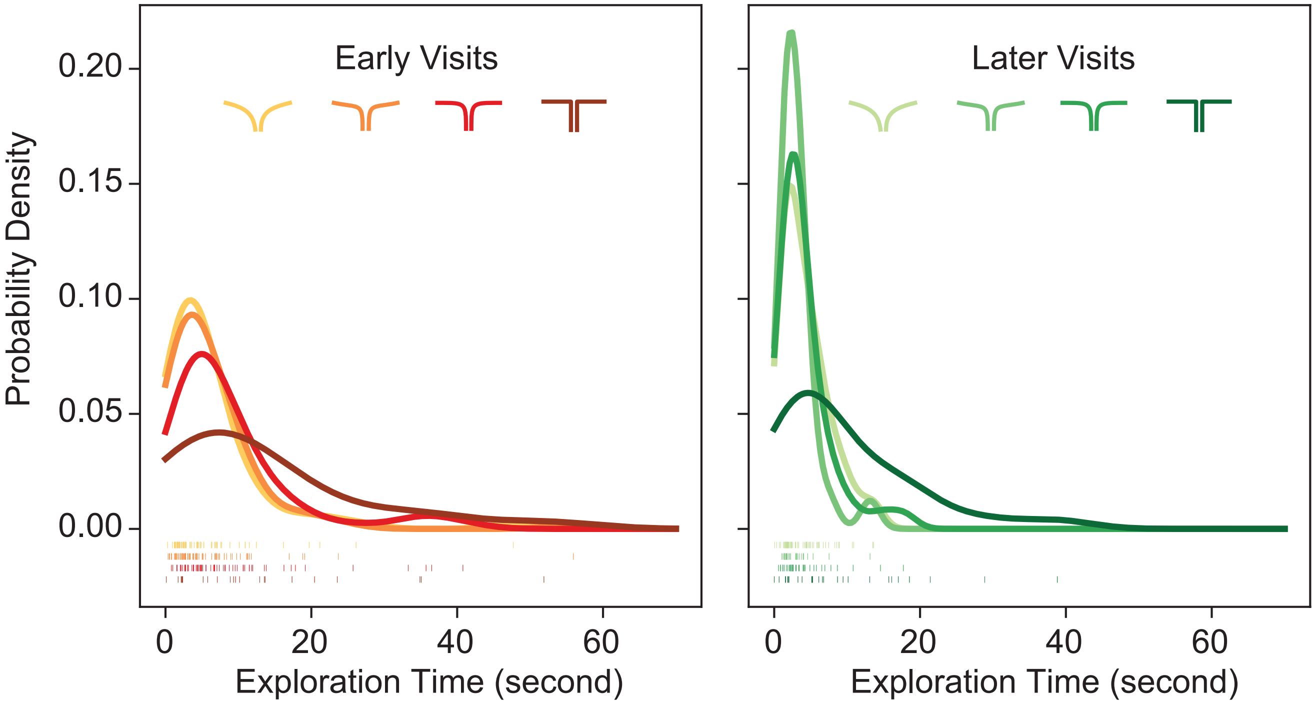


**Fig. 2.** The probability density estimations for early (1–3 visits, left) and later (20–30 visits, right) visits from Fig 2 in the main MS are overlaid on top of each other for each floral shape represented by different hues. (Left) Early visits were similar across all flower shapes (KS test $p > 0.05$ for all pairs). However, when we compute how much the distributions diverge from each other using Kullback–Leibler (KL) divergence, we see a flower shape dependent pattern (KL increases as the floral shape diverges: $0.053$ funnel/near-funnel < $0.109$ funnel/near-flat < $0.228$ funnel/flat flowers) (Right) After learning, however, the exploration times did not differ among all shapes except the flat flower (KS test, $p = 0.97$ for funnel/near-funnel, $p = 0.39$ funnel/near-flat, $p = 0.11$ near-funnel/near-flat, and $p = 1.32e{-}10$ for flat/funnel, $p = 2.67e{-}11$ flat/near-funnel, and $p = 4.05e{-}8$ flat/near-flat)

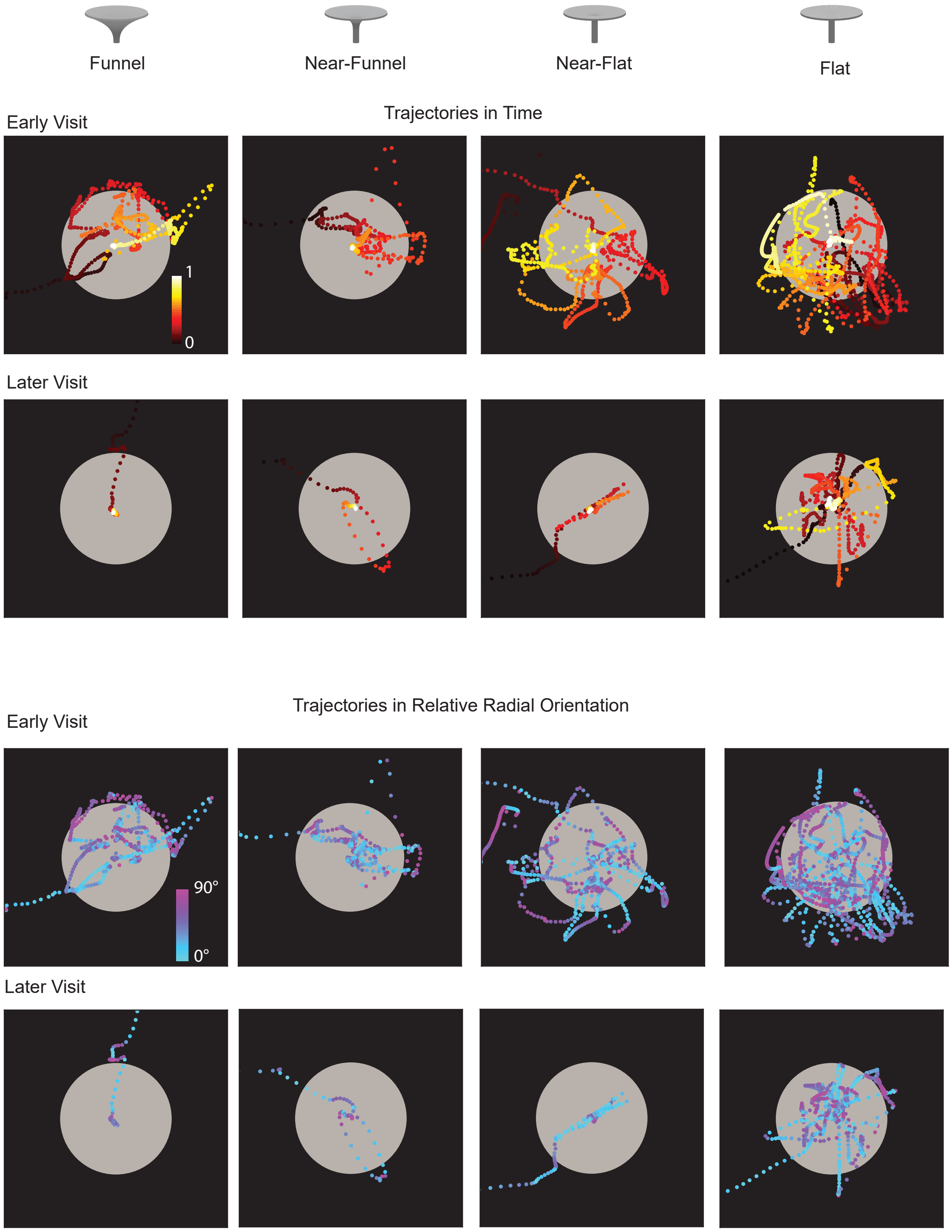


**Fig. 3.** Representative proboscis tip trajectories for a single moth visit are color coded in time (top) and relative radial orientation (bottom) across the four floral shapes. The top row for both color codes represent the first visit a the moth. The bottom row for both color codes each represent the last visit for a moth. The early visit trajectories are the same as the ones in Figure 3 of the main MS.

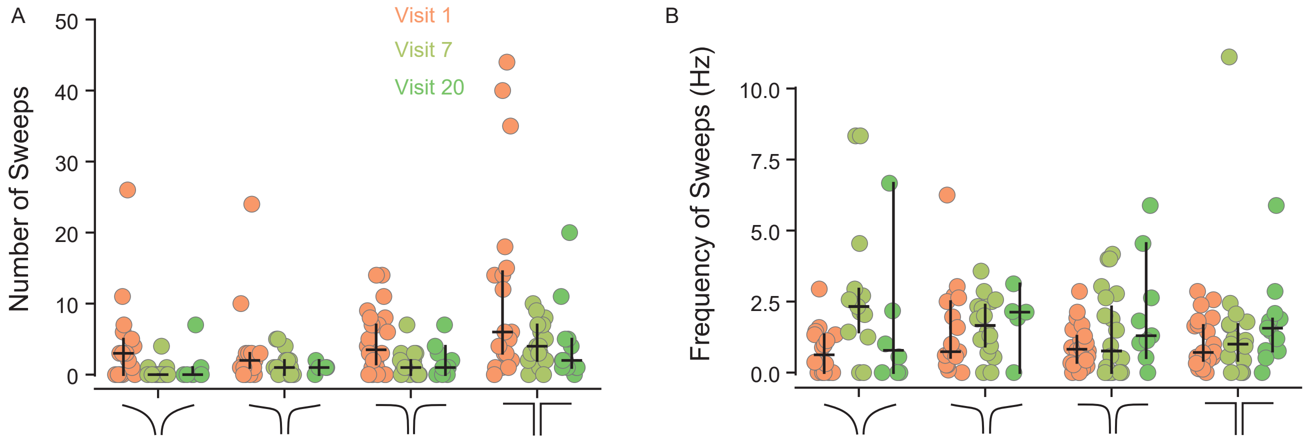


**Fig. 4.** The number of sweeps (A) and the frequency of sweeps (B) are shown for all moths across the different floral shapes for the first visit (brown), the seventh visit (light green) and the twentieth visit (dark green). Each dot represent a single visit for an individual moth. N = Visit 1 - Funnel: 16, Near-Funnel: 17, Near-Flat: 22, Flat: 20, Visit 7 - Funnel: 13, Near-Funnel: 16, Near-Flat: 20, Flat: 17, Visit 20 - Funnel: 8, Near-Funnel: 5, Near-Flat: 9, Flat: 13 moths. The following outliers have not been shown in plot to ensure clarity: Number of Sweeps- 83 (visit 1, Flat flower) and 55 (visit 1, Near-Funnel flower) and Frequency of Sweeps - 25Hz (Visit 20, Funnel Flower)

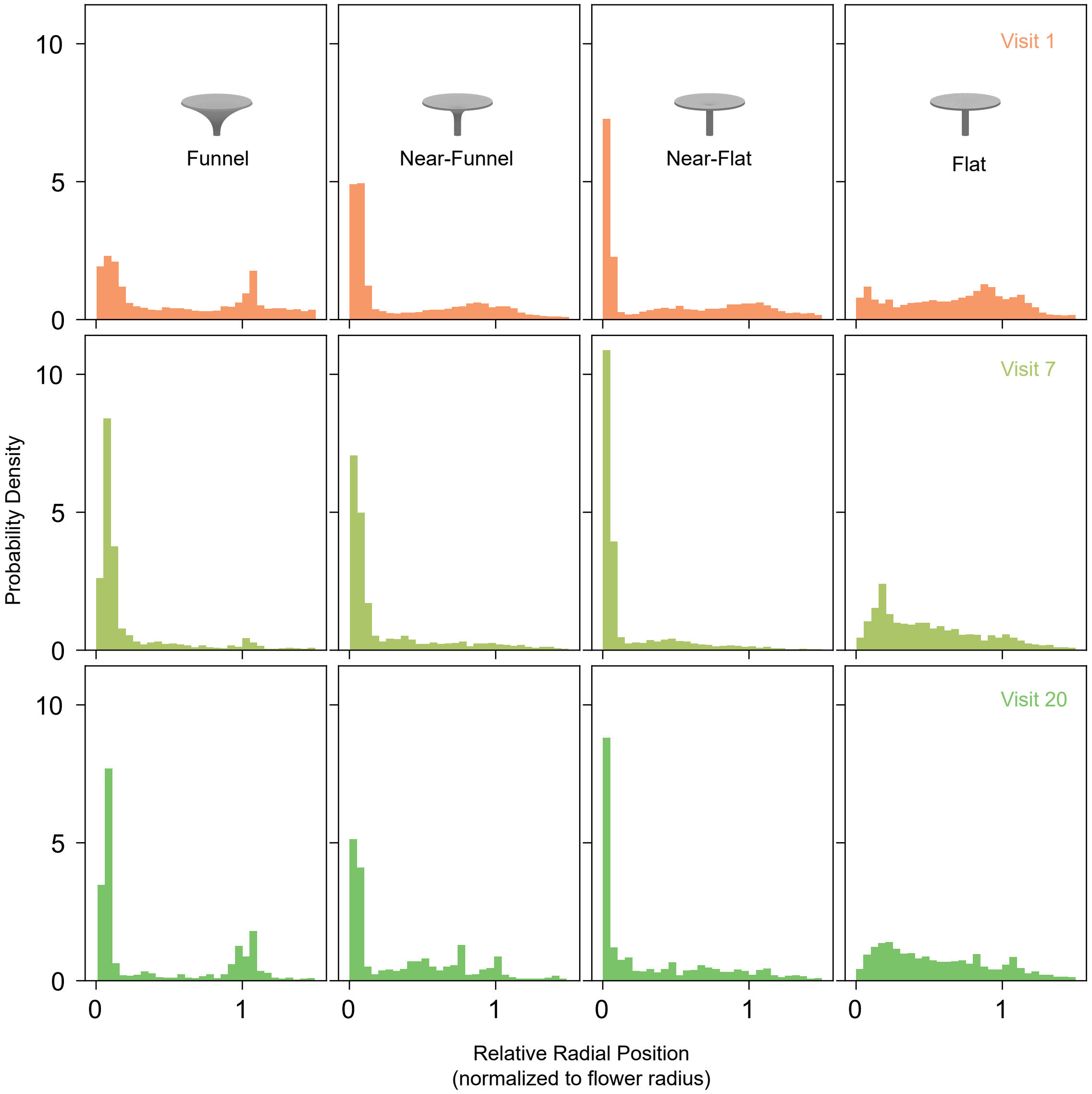


**Fig. 5.** Probability density distributions of the relative radial position of proboscis tip across all moths for the first visit (brown), seventh visit (light green) and twentieth visit(dark green) are shown for the four different flower shapes.

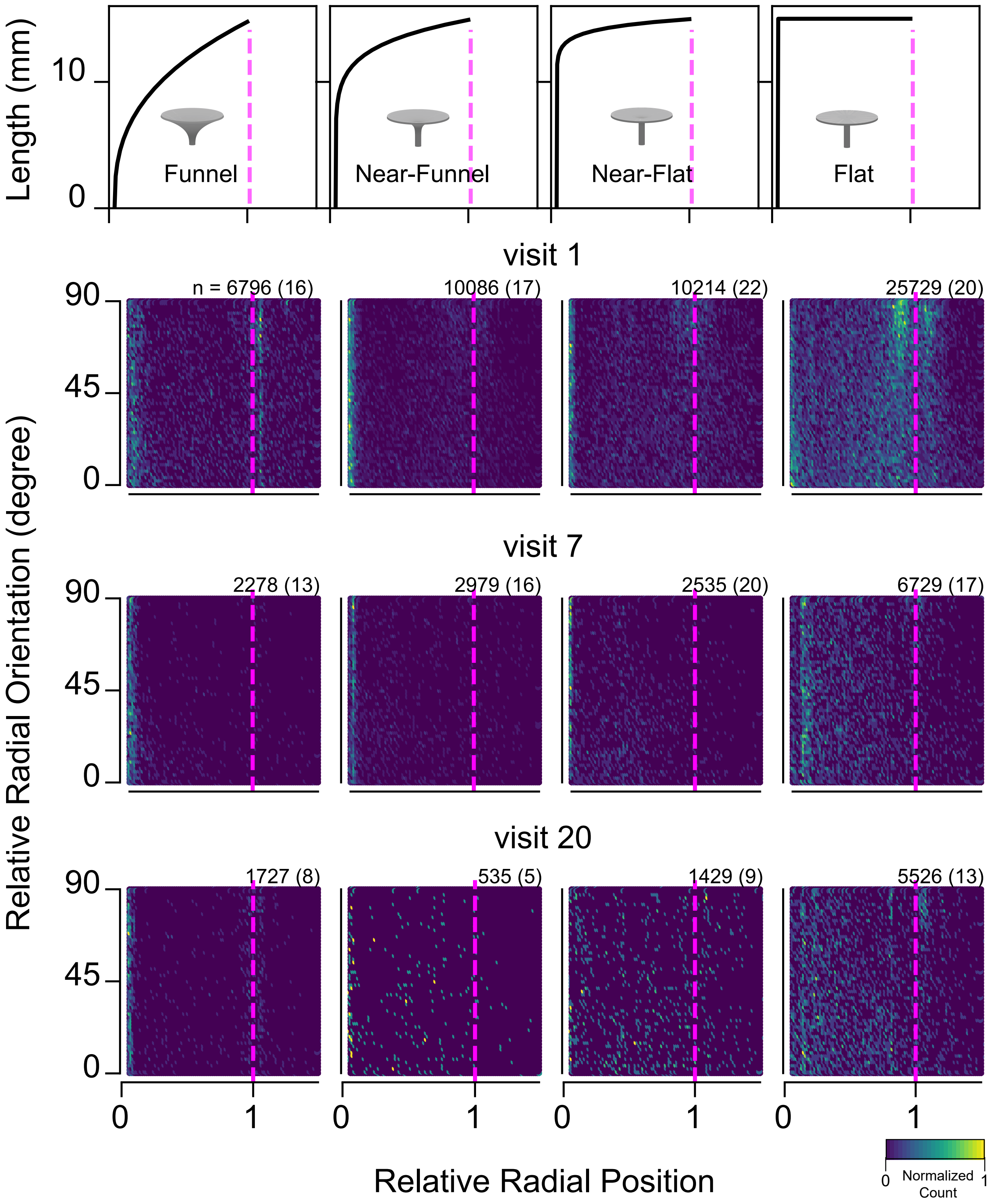


**Fig. 6.** A heat map of the raw data (ploted as hexbins) are shown for the relative radial position and relative radial orientation across of the proboscis tip. These data a shown for the four floral shapes (along the column) and over repeated visits (along the row) polled across all moths. The number on the top left corner of each plot represents the number of frames pooled across the number of moths written in brackets.

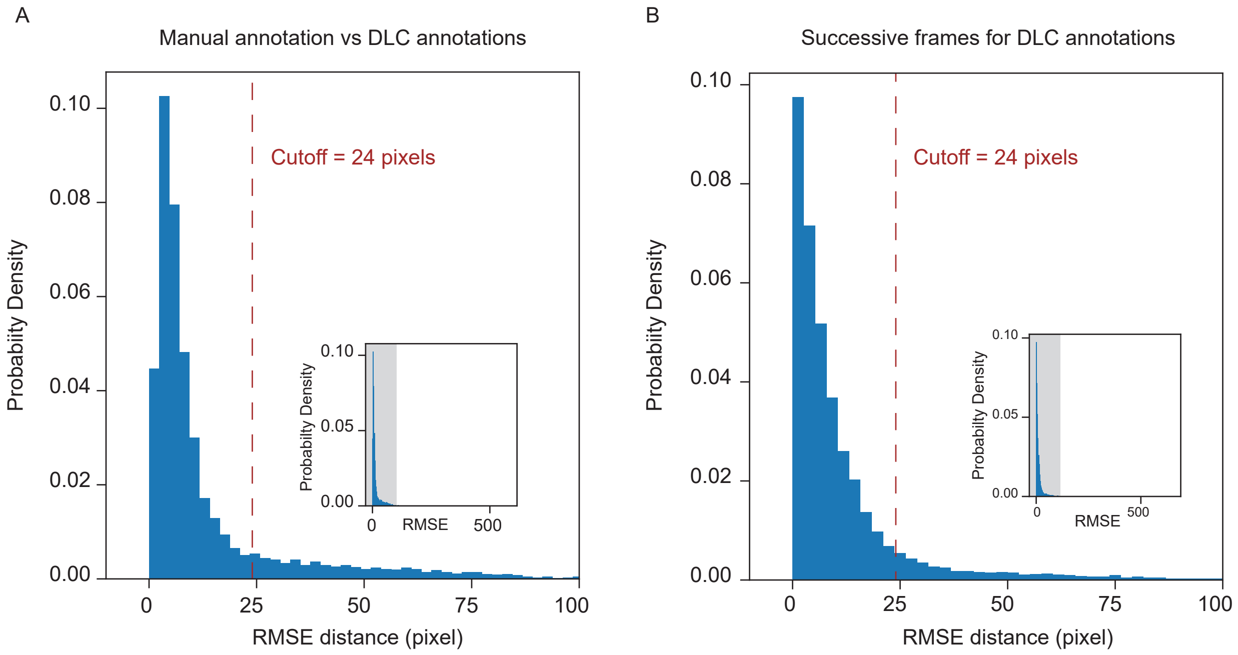


**Fig. 7.** The probability density distribution for the root mean square error (RMSE) distance between the proboscis tip in A) manually annotated versus DeepLabCut (DLC) annotations and in B) successive frames for the DLC annotations are shown here. We manually annotated 6 videos comprising of 10308 frames. Based on the error comparing manual to DLC annotations (shown in A), we used a cut off of 24 pixels. If the distance between the proboscis tip on successive frames with DLC annotations (shown in B) was larger than this cutoff, the tracking data for that frame was dropped. With this cutoff, we could include 87.84% of our data. Inset shows the entire distribution with the grey bar highlighting the region that is zoomed in to visualize the cutoff pixel distance.
